# Induction of Moderate DNA Damage Enhances Megakaryopoiesis and Platelet Production

**DOI:** 10.1101/2025.05.08.652525

**Authors:** Roelof H Bekendam, Virginia Camacho, Andrew P Stone, Maria N Barrachina, Siobhan Branfield, Estelle Carminita, Isabelle C Becker, Dong H Lee, Ethan Walsey, Clementine Payne, Jakub Kaplan, Julia Tilburg, Sharmistha Pal, Luis Francisco Zirnberger Batista, Joseph E Italiano, Kellie R Machlus

**Affiliations:** Vascular Biology Program, Boston Children’s Hospital, Boston, Massachusetts; Department of Surgery, Harvard Medical School, Boston, Massachusetts; Division of Hematology and Hematologic Malignancies, Beth Israel Deaconess Medical Center, Boston, Massachusetts; Biology Department, Brandeis University, Waltham, Massachusetts; Department of Radiation Oncology, Dana-Farber Cancer Institute, Boston, Massachusetts; Department of Medicine, Washington University School of Medicine, St. Louis, Missouri

## Abstract

A common side effect of poly-ADP ribose polymerase (PARP) inhibitors is low platelet counts, or thrombocytopenia, presumably mediated through platelet progenitors, megakaryocytes (MKs). MKs are large, hematopoietic cells with a polyploid, multi-lobulated nucleus. While DNA replication in MKs (endomitosis) is well studied, limited investigations have examined the impact of DNA damage and repair inhibition on megakaryopoiesis. To explore PARP inhibitor-induced thrombocytopenia, we treated mice with PARP inhibitors (niraparib and olaparib), which are approved for the treatment of solid tumors. While high-dose niraparib treatment led to thrombocytopenia, consistent with clinical observations, treatment at a lower dosage led to a significant, >1.5-fold increase in both the number of bone marrow MKs and circulating platelets. This increase was accompanied by elevated DNA damage in both MKs and MK progenitors, as measured by both γH2AX accumulation and comet assays of MKs. Notably, platelets from niraparib-treated mice were functionally normal in their response to ADP, TRAP, and collagen. Gamma-irradiation treatment similarly increased MK and platelet counts in mice, suggesting that moderate DNA damage enhances megakaryopoiesis and increases platelet counts. These data reveal a previously unknown relationship between MKs and DNA damage and present a novel target for triggering enhanced platelet production in vivo.

**Key Points:**

1. Treatment of mice with low dose PARP inhibitors or gamma-irradiation enhances platelet counts.
2. Low dose PARP inhibitor treatment leads to increased DNA damage in MKs and MK progenitors and enhances bone marrow megakaryopoiesis.

## Introduction

DNA is susceptible to damage arising from normal metabolic processes or exposure to DNA damaging agents. The cellular DNA damage response (**DDR**) relies on a complex interplay between pathways that modulate cell survival or cell death.^1^ While generally associated with deleterious responses, accumulation of DNA damage also creates vulnerabilities that are clinically exploited.^2,3^ Therapeutically targeting the DDR is particularly common for the treatment of solid tumors.^4^ Inhibitors of poly-ADP ribose polymerase (**PARP**), a sensor of DNA damage central to the DDR, were first released in 2014.^2^ In physiological conditions, PARP binds to sites of DNA damage and recruits DNA repair effectors that repair DNA lesions. PARP inhibitors (**PARPi**) such as Olaparib and Niraparib “trap” PARP at lesions, leading to DNA replication fork stalling and accumulation of DNA damage through the formation of toxic double-stranded breaks.^2,5^ Notably, PARPi can cause thrombocytopenia in patients,^2,5^ presumably due to deleterious effects of PARP inhibition in platelet-producing cells, megakaryocytes (**MKs**), since platelets lack DNA. MKs are large, polyploid, hematopoietic cells that primarily reside in the bone marrow. MKs can reach states of polyploidy of up to 128N, and contain multiple nuclear lobes within a single nuclear membrane.^6,7^ However, our knowledge of how MKs respond to acquiring many-fold more DNA than other mature blood cells has lagged. In this study, we reveal that induction of moderate DNA damage through treatment with low dose PARPi and irradiation enhanced megakaryo- and thrombopoiesis, presenting a novel trigger of platelet production in vivo.

## Methods

For detailed methodology, see Supplemental Materials.

## Results and Discussion

A common side effect of PARPi treatment in patients is thrombocytopenia.^2,5^ Little is known about the molecular mechanisms of this adverse effect, and overall, about the influence of genetic instability in MK biology. We first aimed to phenocopy PARPi-induced thrombocytopenia by treating mice with different doses of niraparib (Fig 1A). Mice treated with 50 mg/kg niraparib exhibited severe weight loss and markedly reduced platelet counts (Fig 1B), mean platelet volume (**MPV**) and immature platelet fraction (**IPF**) (Supplemental [**S**] Fig 1A, B) after 6 days, which led to premature humane experimental termination. Unexpectedly, however, treating mice with lower dose niraparib (25 mg/kg) led to a significant increase in platelet counts beginning 7 days after treatment induction and persisting through the end of the experiment (day 11) (Fig 1C). These results suggest that there is a threshold response at which niraparib displays opposing effects on platelet production, consistent with preclinical rat studies.^8,9^ To substantiate this, we measured MPV (Fig 1D) and IPF (Fig 1E); both were elevated in mice treated with low dose niraparib, suggestive of new platelet production. Additionally, thiazole orange (**TO**) staining of platelets on day 11 revealed that mice treated with low dose niraparib had a higher proportion of reticulated platelets (Fig 1F), indicative of young platelets age in circulation.^10^ Upon sacrifice, the spleen weights were similar between mice treated with vehicle or 25 mg/kg niraparib for either 3 or 11 days, suggesting no difference in platelet mobilization to or from the spleen (Fig 1G). Quantification of peripheral blood cell counts revealed that mice treated with 25 mg/kg niraparib for 11 days had a concurrent decrease in hemoglobin, but no significant change in white blood cell count (Fig S1C, D). Notably, a similar increase in platelet count was observed when mice were treated with the structurally unrelated PARPi compound olaparib, indicating that this effect was not niraparib-specific (Fig S1E, F).

**Figure 1.**
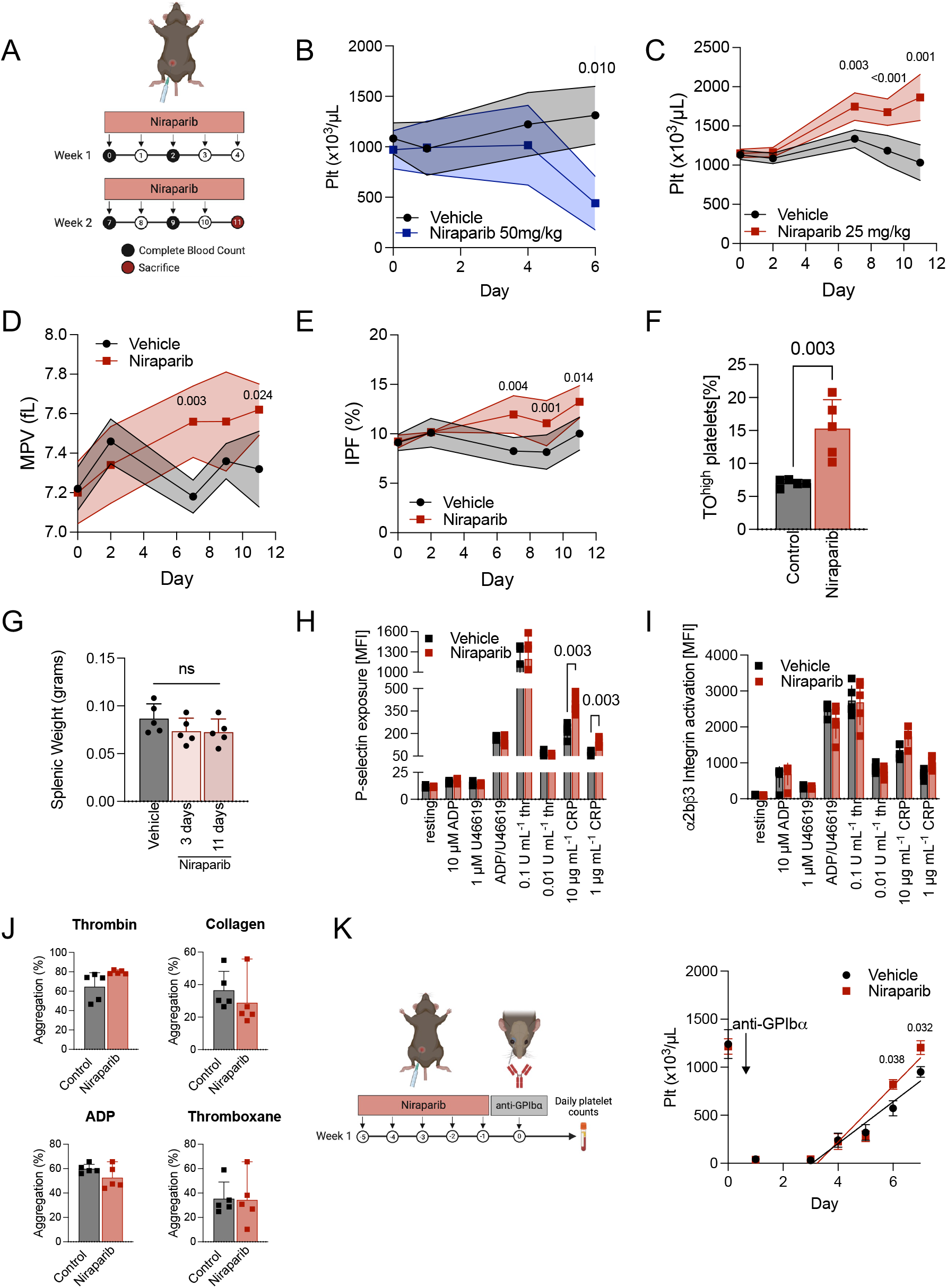
Low dose Niraparib treatment enhances platelet count in mice without affecting platelet function. **A)** Schematic outlining niraparib administration: Mice were injected intraperitoneally with niraparib or vehicle (DMSO) at the indicated times. Blood was drawn via tail vein venipuncture and complete blood counts (CBC) were measured using a Sysmex hematology analyzer. **B)** Platelet counts in vehicle or niraparib-treated mice (50 mg/kg); n=5 mice/group. **C)** Platelet counts in vehicle or niraparib-treated mice (25 mg/kg); n=5 mice/group. **D)** Mean platelet volume (MPV) in vehicle- or niraparib-treated mice (25 mg/kg); n=5 mice/group. **E)** Immature platelet fraction (IPF) in mice treated with niraparib (25 mg/kg); n=5 mice/group. Data in B) - E) shown as mean ± SD. Two-way ANOVA with Tukey’s correction for multiple comparisons. **F)** Frequency of reticulated, thiazole orange (TO)^high^ platelets in whole blood of vehicle- or niraparib-treated mice (25 mg/kg); n=5 mice/ group. Data as mean ± SD, unpaired, two-tailed student’s t-test. **G)** Spleen weights from vehicle- or niraparib-treated mice (25 mg/kg) after 3 or 11 days of treatment, as indicated; n=5 mice/ group. Data as mean ± SD, unpaired, one-way ANOVA with Šidák correction for multiple comparisons. **H)** Surface P-selectin exposure and **I)** Activated αIIbβ3 integrins on platelets in whole blood were measured by flow cytometry. Mean fluorescence intensity (MFI; geometric mean) is shown; n=5 mice/group. Data as mean ± SD, multiple t-tests. **J)** Platelet aggregation was measured using washed platelets isolated from vehicle- and niraparib-treated mice. Absorbance at 450 nm was used to calculate total aggregation for each sample normalized to the positive and negative controls; n=5 mice/ group. Data as mean ± SD, unpaired two-tailed student’s t-test. **K)** Schematic of in vivo platelet depletion (left) using an anti-GPIbα antibody (2 µg/g per mouse). Platelet counts following anti-GPIbα administration were measured over 6 consecutive days; n=5 mice/ group. Data as mean ± SD, unpaired, two-tailed student’s t-test.

We next tested platelet functionality in 25 mg/kg niraparib- and vehicle-treated mice. Baseline expression of αIIb integrins (**CD41**), glycoprotein (**GP**)Ibα, and GPVI were unaltered (Fig S1G), as was platelet activation when whole blood was stimulated with ADP and thrombin (Fig 1H, I). We did, however, observe heightened sensitivity to GPVI stimulation in platelets from niraparib-treated mice treated with collagen-related peptide (**CRP**). This may be attributed to the increase in reticulated platelets, which often display increased GPVI reactivity (Fig 1H, I).^11,12^ However, platelets from niraparib-treated mice had unaltered aggregation, suggesting that transient low-dose niraparib treatment does not elevate thrombotic risk (Fig 1J).

Finally, to evaluate the impact of 25 mg/kg PARP inhibition on platelet recovery after thrombocytopenia, we preconditioned mice with niraparib, followed by platelet depletion using an anti-GPIbα antibody. Mice preconditioned with niraparib showed faster recovery and higher platelet counts (Fig 1K) after induction of thrombocytopenia. Together, these findings demonstrate that treatment with 25 mg/kg niraparib enhanced the production of functional platelets in mice.

To probe the etiology of niraparib-induced platelet production, we assessed the effect of niraparib on hematopoietic stem and progenitor cell (**HSPC**) populations (gating strategy outlined in Fig S2). We observed an increase in multipotent progenitor populations 2 (**MPP2**) and 3 (**MPP3**) after 3 days of treatment. After 11 days of treatment, we observed a significant increase in HSCs, MK progenitors (**MkPs**), and MKs, in addition to all MPP populations, while CD41^+^ HSCs were unaltered (Fig 2A). These data suggest that niraparib treatment primes HSC differentiation towards the MK lineage. This was validated by immunofluorescent visualization of MKs in femoral cryosections (Fig 2B), revealing an increase in both the number (Fig 2C), and size (Fig 2D) of MKs in niraparib-treated mice after 11 days. To confirm these findings in vivo, we examined MKs via 2-photon intravital microscopy (**2PIVM**) in the calvaria of MK/platelet reporter mice (von Willebrand factor (**vWF**)-GFP).^13^ In vivo MK quantification and phenotyping also showed a significant increase in MK numbers upon niraparib treatment, corroborating our findings from the in situ analysis and confirming that MKs were indeed making proplatelets (Fig 2E, F; Supplemental movies S1, 2). To further investigate MK morphology, we performed electron microscopy (**EM**) on femoral whole bone marrow sections from both vehicle- and niraparib-treated mice (Fig S3). EM images revealed no gross changes or abnormalities in MK structure, bone marrow cellularity, or the endothelial cells lining vascular lumens (representative images, Fig S3).

**Figure 2.**
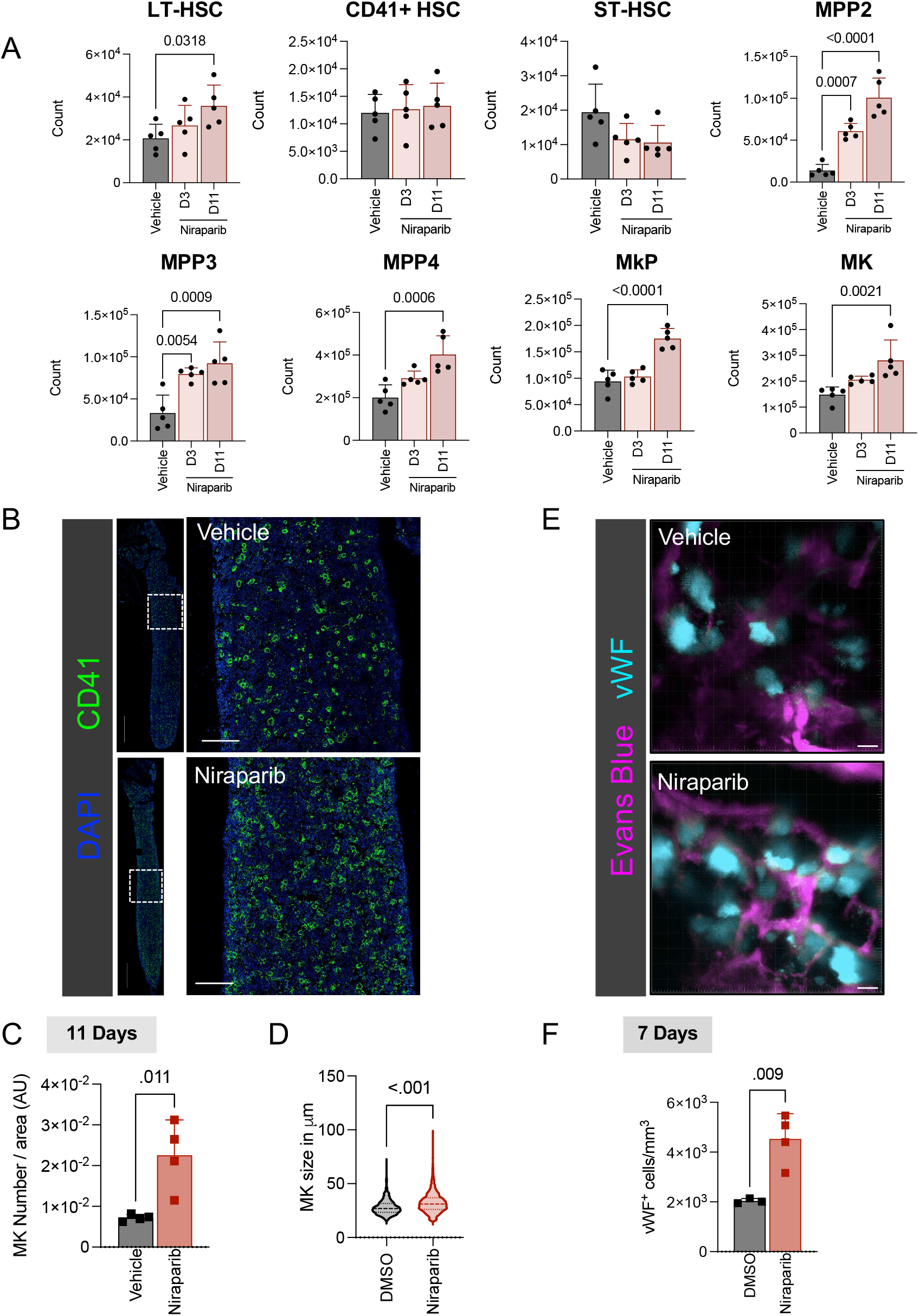
Low dose Niraparib treatment enhances megakaryopoiesis. **A)** HSPC counts in vehicle- or niraparib-treated mice; n = 5 mice/ group. Data shown as mean ± SD, One-way ANOVA with Dunnett’s post hoc test. Gating strategy shown in Supplemental Figure 2. **B)** Representative images of femoral cryosections from vehicle or niraparib treated mice (CD41, green). Nuclei were counterstained using DAPI (blue). Scale bars: 1000 µm; inset scale bars: 200 µm. **C)** MK counts and **D)** area in vehicle- or niraparib-treated mice were quantified in femoral cryosections; n=4 mice/group. Data shown as mean ± SD, unpaired, two-tailed student’s t-test. **E)** MK/platelet reporter mice (vWF-GFP) were treated with 25 mg/kg niraparib or vehicle for 7 days and 2-photon intravital microscopy of the calvaria was performed. Representative images show MKs (cyan) and vasculature (magenta, labelled using Evans Blue). Scale bars: 20 µm. **F)** Live MKs were quantified by size and GFP signal; n=4 mice/group. Data shown as mean ±?SD, unpaired, two-tailed student’s t-test.

Notably, one course of niraparib treatment significantly increased platelet counts for over two weeks (Fig S4A). However, after approximately 6 weeks there were no significant differences in HSPC counts (Fig S4B) or platelet aggregation (Fig S4C) and activation (Fig S4D), suggesting that bone marrow cells can recover from short-term niraparib treatment.

As PARP inhibition impacts DNA repair,^14^ we next aimed to examine DNA damage accumulation in HSPCs following niraparib treatment. To probe the extent and specificity of niraparib-induced DNA damage, we quantified accumulation of the DNA damage marker gamma H2A histone family member X (γ**H2AX**) in HSCs and MkPs. Notably, after three days of Niraparib treatment, and preceding the increase in platelet counts, there was an increase in γH2AX-positive HSCs, CD41^+^ HSCs, and MkPs (Fig 3A).

**Figure 3.**
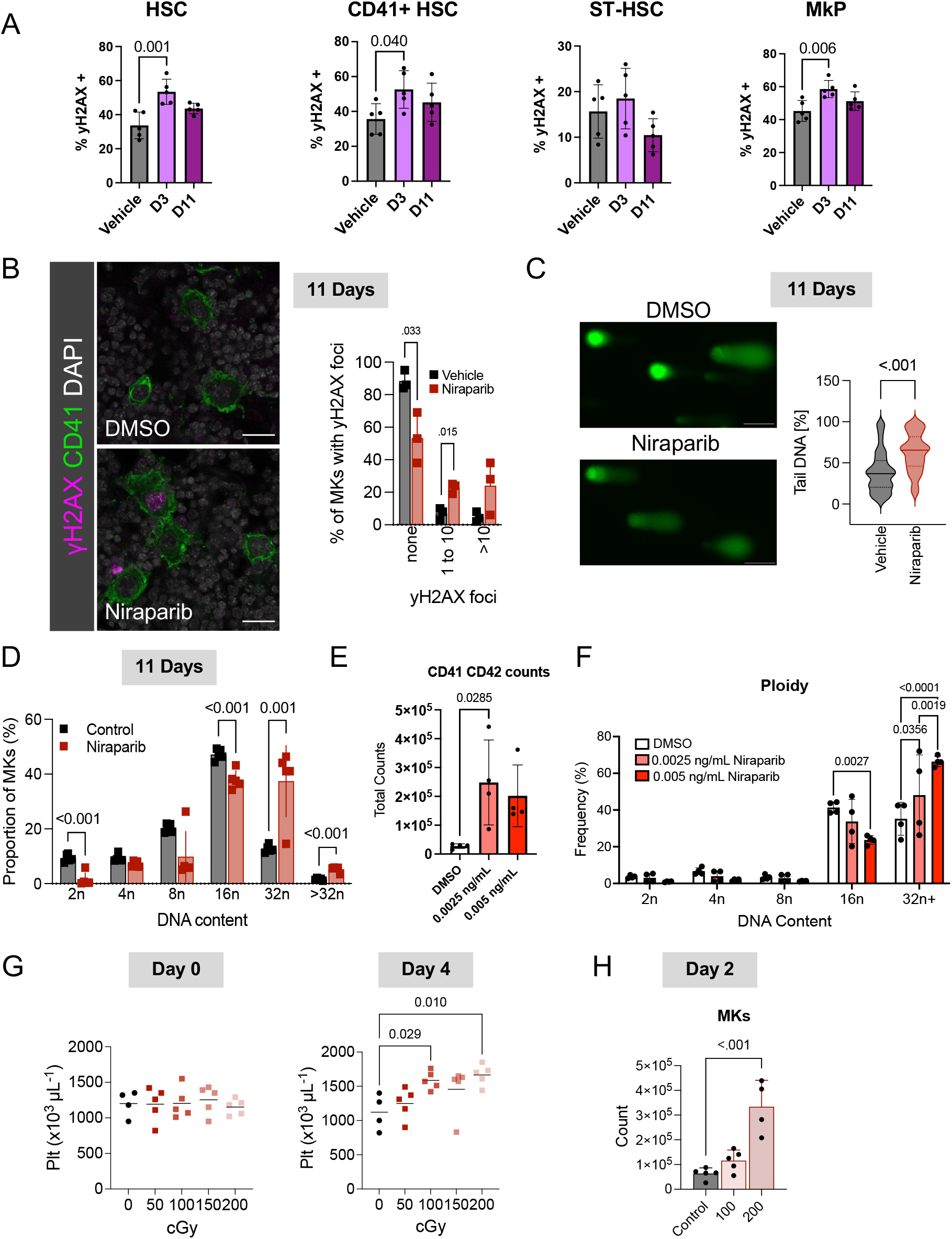
DNA damage dose dependently enhances megakaryopoiesis in vitro and in vivo. **A)** yH2AX positivity in indicated HSPC populations in vehicle- or niraparib-treated mice; n=5 mice/group. Data shown as mean ± SD, One-way ANOVA with Dunnett’s post hoc test. Gating strategy shown in Supplemental Figure 2. **B)** Representative widefield images of femoral cryosections derived from vehicle- or Niraparib-treated mice; (CD41; green), (γH2AX; purple). Scale bars: 20 µm. Quantification shows the percentage of femoral MKs that have either 0, 1-10, or >10 γH2AX foci in the nucleus in vehicle or niraparib treated mice; 100 MKs counted per femur, n=3 mice/group. Data shown as mean ± SD, multiple t-tests. **C)** Representative images of native MKs from vehicle- or Niraparib-treated mice (left); (green =SYBR Gold). Scale bars: 100 µm. MKs were isolated from the femurs of vehicle- or niraparib-treated mice. The tail DNA (%) of at least 120 MKs per femur were analyzed across three replicates (3 femurs) per condition; n=3 mice/group. Violin plots show distribution of all data points and median and interquartile range, unpaired, two-tailed student’s t-test. **D)** Ploidy distribution in native MKs from vehicle- or niraparib-treated mice; n=5 mice/group. Data shown as mean ± SD, Two-way ANOVA with Šidák correction for multiple comparisons. **E)** Number of MKs (CD41/CD42d^+^) and **F)** ploidy distribution was assessed after treatment of HSPCs with indicated doses of niraparib in vitro; n=5 mice/group. Data shown as mean ± SD, One-way ANOVA with Dunnett’s post hoc test. **G)** Platelet counts at day 0 (left) and day 4 (right) following sublethal irradiation; n=4-5 mice/dose. Data shown as mean ± SD, unpaired, One-way ANOVA with Tukey’s correction for multiple comparisons. **H)** Murine bone marrow MKs (CD41/CD42d^+^) were quantified by flow cytometry 2 days after treatment with 0, 100, or 200 cGY; n=4-5 mice/group. Data shown as mean ± SD, unpaired, One-way ANOVA with Šidák correction for multiple comparisons.

Few studies have directly addressed the role of DNA damage in MKs. However, MK development has been associated with DNA damage and repair,^15,16^ suggesting a connection between these processes. Further, DNA damage of committed HSCs can promote MK differentiation via arrest in the G2 phase of the cell cycle.^17^ Probing γH2AX immunofluorescence in situ, we observed a higher proportion of γH2AX positive MKs and more foci per MK (Fig 3B) after niraparib treatment. To directly analyze DNA breaks in native MKs, we next isolated bone marrow MKs from niraparib- and vehicle-treated mice and performed comet assays (Fig 3C). These assays showed significantly longer tail lengths, indicating increased DNA damage, in MKs from niraparib-treated mice (Fig 3C). Together, our data demonstrate that the lower niraparib concentration utilized in our assays was sufficient to induce DNA damage in MKs.

In addition to changes in MK numbers, niraparib treatment also resulted in striking differences in MK ploidy from day 3 to day 11 (Fig S4E and 3D), with a shift towards high-ploidy MKs. These data further substantiate that despite increased DNA damage accumulation, MKs continued to mature and replicate DNA. While MK ploidy is not directly causative of platelet production, higher ploidy MKs with more cytoplasm are thought to largely contribute to platelet production.^18^ Collectively, these data indicate that inducing low levels of DNA damage in MKs promoted megakaryopoiesis resulting in increased platelet production.

To rule out effects from secondary mediators in vivo, we performed in vitro experiments in which murine bone marrow HSPCs^19,20^ were treated with niraparib and MK differentiation was assessed after 4 days in culture. Niraparib treatment resulted in significantly more HSPCs differentiating into (CD41/CD42d^+^) MKs (Fig 3E) without a change in viability (Fig S4F). Further, consistent with in vivo data, the increase in MKs was accompanied by a shift towards higher ploidy (Fig 3F).

Finally, to confirm that the increase in MK and platelet counts was not specific to PARPi, but rather a general response to controlled DNA damage, we subjected mice to a dose curve of ionizing radiation. Radiation dose dependently increased platelet counts after 4 days (Fig 3G). Similar to niraparib, 2 days post-irradiation, preceding the increase in platelet counts, bone marrow MKs were also significantly increased in a dose-dependent manner (Fig 3H), accompanied by an accumulation of DNA damage in the MKs of mice irradiated with 200cGy.

In summary, our results reveal that moderate DNA damage acts as a potent driver of megakaryopoiesis, polyploidization, and subsequent platelet production in vivo. These data uncover a previously understudied relationship between MKs and DNA damage and present a novel mechanism governing platelet production in vivo.

## Acknowledgements

This work was supported by the National Institute of Diabetes and Digestive and Kidney Diseases (R03DK124746 to KRM) and the National Heart, Lung, and Blood Institute (R01HL151494 to KRM K99HL175037 to MNB and R35HL161175 to JEI) at the National Institutes of Health; RHB is the recipient of a 2023 HTRS Mentored Research Award from the Hemostasis and Thrombosis Research Society (HTRS), supported by and educational grant from CSL Behring. MNB is the recipient of fellowships from the American Society of Hematology (ASH Restart Award) and the American Heart Association (23POST1011433). KRM is the recipient of an Innovative Project Award from the American Heart Association (24IPA1274573). VC is the recipient of an ASH Scholar Award. SP is supported by funds from U54 grant 1U54CA274516 and NCI SPORE 2P50CA165962. LFZB was supported by the National Institutes of Health (CA258386; HL174789; HL172961), the Department of Defense (BM230053), the American Cancer Society (133856-RSG), and the Siteman Cancer Center at Washington University in St. Louis.

## Authorship Contributions

RHB, APS, CP, VC, EC, SB, MNB, ICB, JK, DHL, EW, and JT performed experiments, interpreted and analyzed data. RHB, VC, APS, CP, MNB, JK, LFZB, SP, JEI and KRM conceptualized the experiments and research ideas. RHB, VC, CP, LFZB, ICB, JEI and KRM wrote and edited the manuscript. SP, JEI, and KRM funded the research.

---

**Figure S1.**
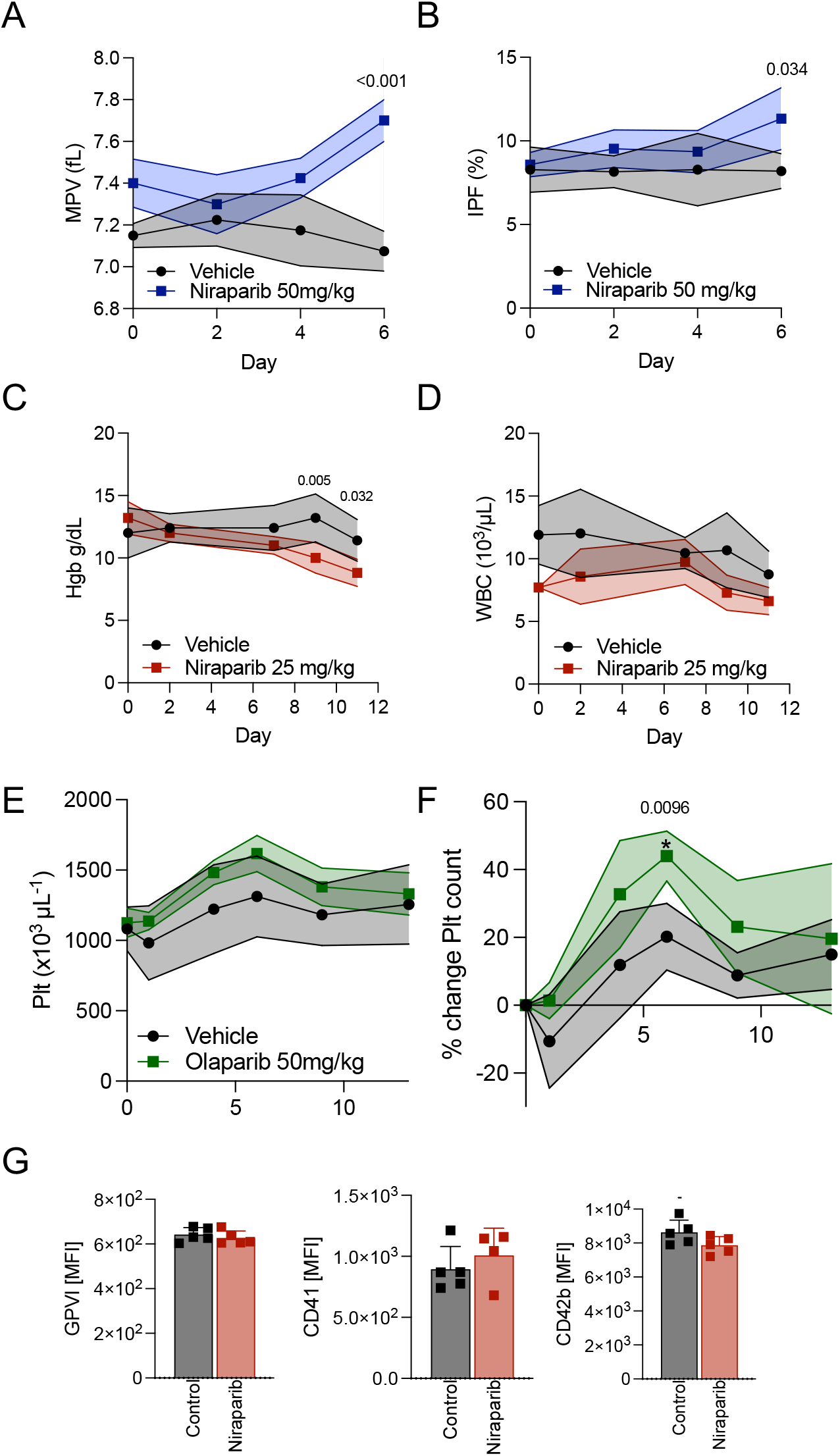

**Figure S2.**
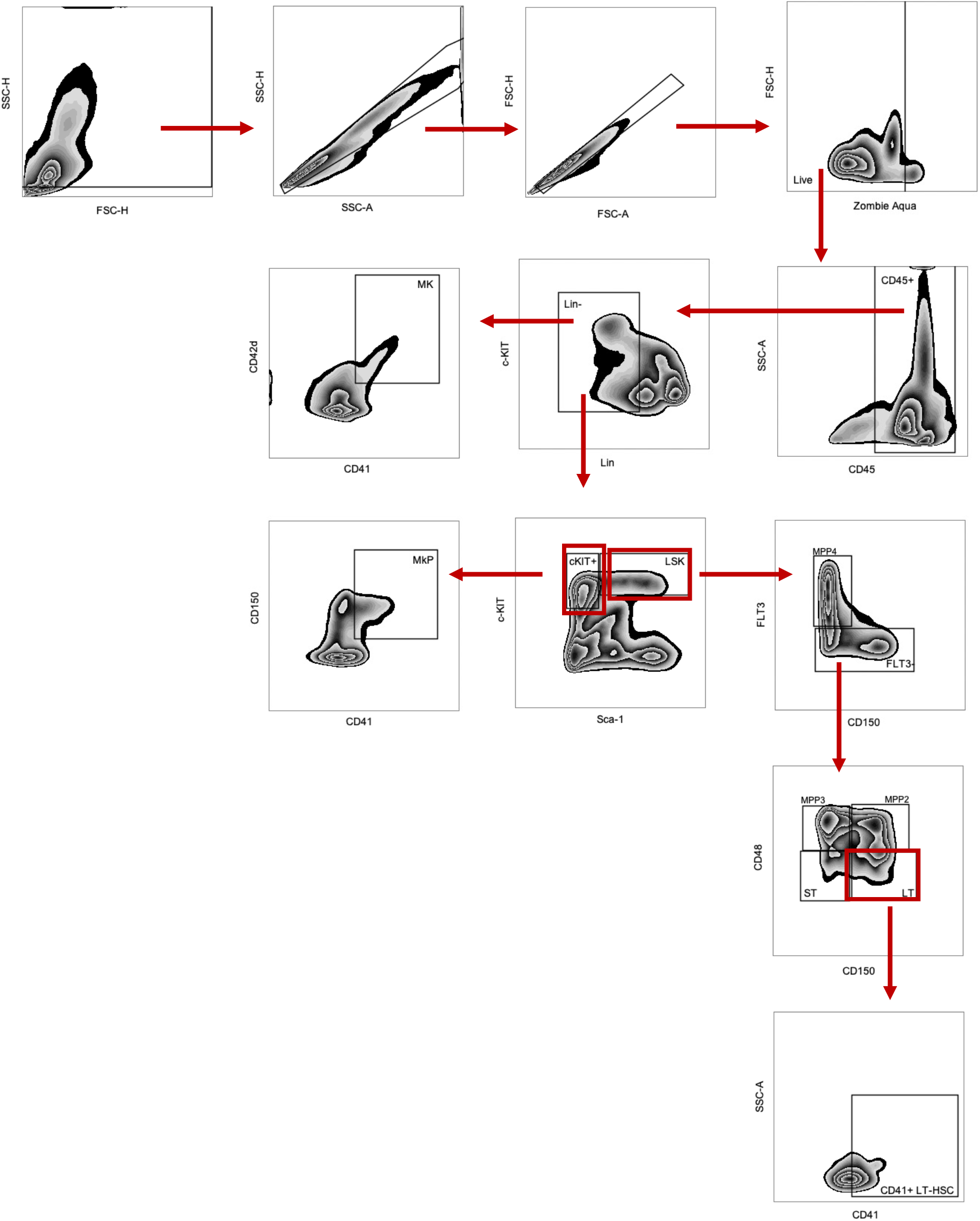

**Figure S3.**
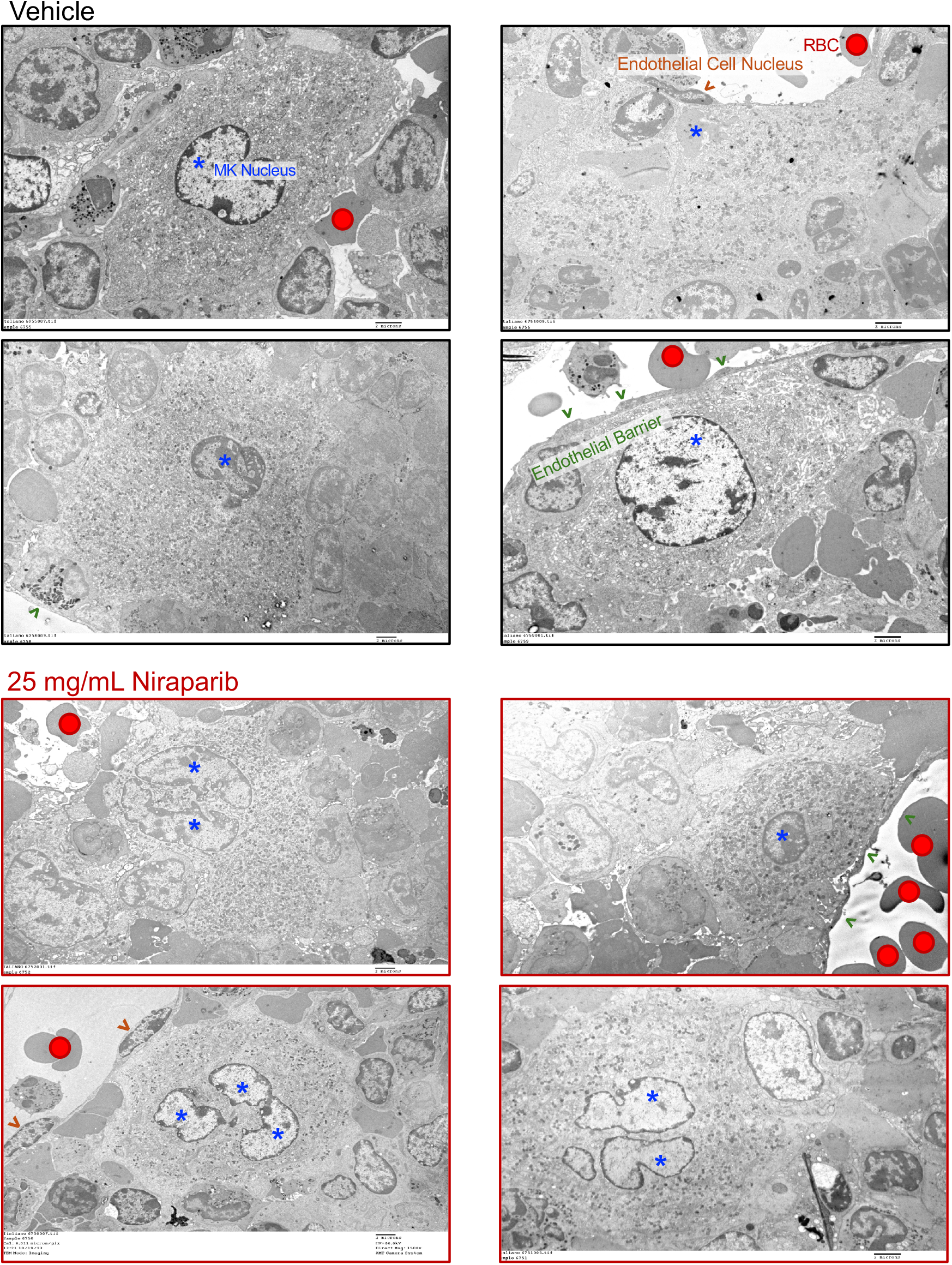

**Figure S4.**
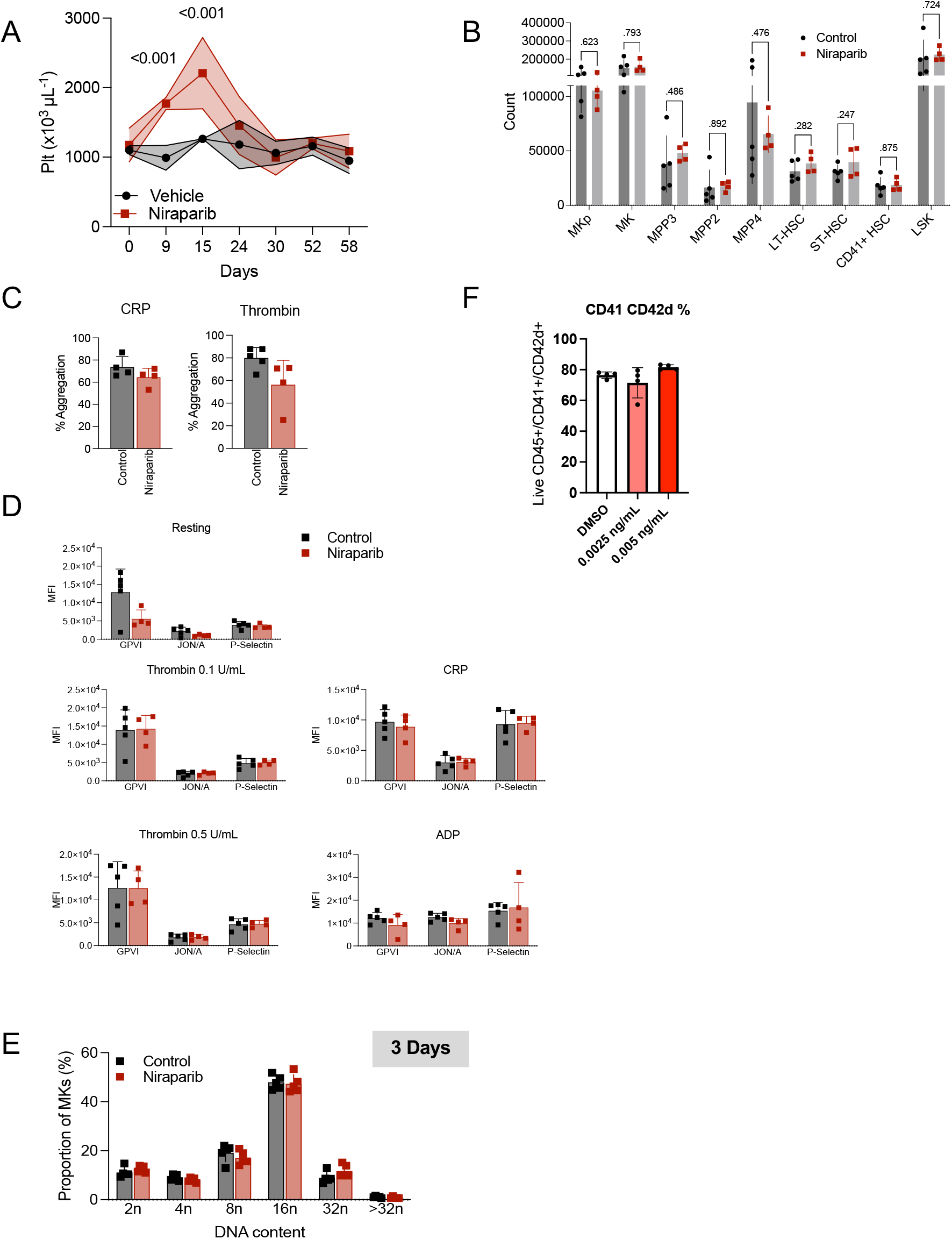

