## Supplemental Methods and Figure Legends for "Induction of Moderate DNA Damage Enhances Megakaryopoiesis and Platelet Production"

**Supplementary Materials**

Supplementary Methods

### Reagents

Niraparib (MK-4827, Sellekchem) was dissolved in 100% DMSO to a stock concentration of 50 mg/mL. For in vivo experimentation stock concentration was diluted with 10% (2-hydroxypropyl)-ß-cyclodextrin (332607, Millipore Sigma) in sterile PBS to reach a final concentration of either 50 mg/kg body weight or 25 mg/kg body weight.

### Mice

C57BL/6J mice (#027C57BL/6) were purchased from Charles River Laboratories (Worcester, MA, USA). All procedures were approved by the international animal care and use committee (IACUC) at Boston Children's Hospital (Protocol 00001248).

### Comet assay

Alkaline Comet Assays were performed using the CometAssay® Kit (34250-50K, Trevigen Inc.) according to the manufacturer's instruction. Briefly, native bone marrow megakaryocytes (MKs) were isolated as previously published,[1] and $1x10^3$ cells were resuspended in LM Agarose and spotted on CometSlide. To solidify, the agarose slides were kept at 4°C for 30 min and then immersed in cold Lysis Solution for 16-18 hours at 4°C. The next day, the slides were immersed in an alkaline unwinding solution (300 mM NaOH and 1 mM EDTA) for 2 hours at 4°C in the dark before the slide was subjected to electrophoresis in alkaline electrophoresis solution (300 mM NaOH, 1 mM EDTA; pH>13) at 32V, 1.48V/cm for 1 hour in the cold room. Subsequently, the slides were immersed in nuclease-free water for 5 min, dehydrated in 70% ethanol for 5 min, and then dried at 37°C for 15 min in the dark. Finally, to visualize DNA, SYBR Gold staining was performed for 30 min at room temperature in the dark before the slides were washed in nuclease-free water and dried at 37°C. Images were taken at 10X using the Zeiss microscope. Comets were analyzed using the OpenComet plugin in the ImageJ software. To determine the % tail, OpenComet software determined the head and tail of each MK. The tail is defined as the elongated smear adjacent to the circular head. All scans were manually evaluated to ensure the correct detection. MKs were isolated from the femurs of three mice per condition and at least 120 MKs were analyzed per femur.

### 2-Photon intravital microscopy

2-photon intravital microscopy (2PIVM) was performed on calvaria of C57BL/6JR (von Willebrand factor [VWF]-eGFP) mice treated with niraparib (25 mg/kg) or vehicle for 7 days. MKs and platelets (enhanced green fluorescent protein [$eGFP^+$]) and vessels (Evans blue) were visualized using a 960 nm laser line. Mice were anesthetized and a window in the calvaria was made to view the exposed bone marrow. MK distribution and proplatelet formation were observed and imaged for 5 minutes in 4 different fields of view per mouse.[2]

**Flow cytometric analysis of murine platelets in whole blood**

After terminal exsanguination into EDTA-coated tubes, whole blood (5 µL) was diluted in 500 µL of Tyrode's buffer (J67607, Alfa Aesar) and stained with thiazole orange (TO; 200 ng/mL) for 15 min. Flow cytometric analysis (Accuri C6 plus, BD Biosciences) was performed to identify the frequency of TO(+) platelets. To measure platelet activation, 50 µL of murine whole blood was washed in Tyrode's Buffer, and indicated doses of adenosine diphosphate (ADP), U46619, thrombin, and cross-linked collagen-related peptide (CRP-XL) were incubated with platelets for 15 mins. P-selectin exposure (WUG 1.9-FITC, Emfret Analytics) and activation of αIIbβ3 integrins (JON/A-PE, Emfret Analytics) were determined by flow cytometry. Cell surface glycoprotein (GP) expression on the surface of unactived platelets was measured in whole blood (CD41: 133904, Biolegend, GPIbα: #M040-1, Emfret Analytics; GPVI: #M011-1, Emfret Analytics).

**Platelet aggregometry with washed platelets**

Mice were terminally bled and whole blood was collected in 3.2% citrate. Whole blood was centrifuged at 100*g* for 10 minutes to isolate platelet rich plasma (PRP). Prostaglandin and apyrase were added to PRP and centrifuged at 900*g* for 10 minutes. Plasma was removed from atop the platelet pellet and the platelets were resuspended in 400 µL of Tyrode's buffer. Washed platelets were quantified using a Sysmex cell counter and normalized to a cell count of 1.5 x $10^5$. Forty µl of platelets was incubated with Thrombin (0.5U/mL) or CRP in the presence of 20 µM calcium in a thermoblock at 37° C and shaken at 1200 rpm for 5 minutes. Platelets without agonist or calcium were used as a negative control and defined as 0% aggregation. Platelets in Tyrode's buffer were used as a positive control; defined as 100% aggregation. Absorbance value at 450nm was used to calculate total aggregation for each sample normalized to the controls upon agonist stimulation. Aggregation was calculated as follows:

1. Total Aggregation in each sample = (100% Aggregation) – (0 % Aggregation).
2. Change of absorbance of sample = (Total Aggregation in each sample - Absorbance value at 450nm of sample).
3. (%) of aggregation = Change of absorbance of sample) / (Total aggregation) x 100.

**Ex vivo MK culture and Niraparib treatment**

MKs were differentiated from murine bone marrow hematopoietic stem and progenitor cells HSPCs, as previously published.[2,3] Briefly, HSPCs were enriched using rat anti-mouse lineage panel (133307, BioLegend), followed by magnetic bead isolation using DynabeadsTM (11415D, ThermoScientific) and cultured with TPO (50 ng mL-1) for 5 days. On day 1 HSPCs were cultured in the presence of Dimethyl sulfoxide (DMSO) or Niraparib. Differentiated MKs were quantified and defined by gating on a Live, $CD45^{+}Lin^{-}$ $CD41^{+}$ $CD42d^{+}$ cells.

**In vivo administration of PARP inhibitors**

For the 11-day treatment, C57BL/6J mice were injected intraperitoneally with either niraparib (25 or 50 mg/kg) or vehicle control for 5 consecutive days, followed by two days off, followed by 4 consecutive days of injections. For the three-day treatment, mice were injected intraperitoneally for 3 consecutive days. Similarly, olaparib 50 mg/kg or vehicle was injected intraperitoneally in C57BL/6J mice for the 13-day treatment for 5 consecutive days, followed by three days off, followed by 5 consecutive days of injections. For non-terminal platelet counts, tail vein venipunctures were performed on specified days. For the long-term monitoring experiment, tail vein punctures continued at indicated days after 11-day treatment cessation. Platelet count, IPF, and MPV were quantified using a Sysmex hematology analyzer.

**Murine model of gamma irradiation**

C57BL/6J mice were irradiated with 0, 50, 100, 150, or 200 cGy (Gammacell 40 extractor). Platelet counts were measured prior to and following irradiation by tail vein venipuncture using a Sysmex hematology analyzer. Bone marrow was isolated as described under 'HSPC and MK quantification by flow cytometry'.

**Induction of thrombocytopenia**

C57BL/6J mice were intraperitoneally injected for 5 days with either niraparib 25mg/kg or vehicle control (day -5; D-5, through day -1; D-1). On day 0 (D0) an anti-GPIbα antibody (R300, 2 μg/g, Emfret Analytics) was administered to induce thrombocytopenia. Platelets were drawn by tail vein venipuncture or venipuncture from the retro-orbital plexus and measured using a Sysmex hematology analyzer prior to and following induction of thrombocytopenia until recovery.

**Femur processing and immunofluorescence**

Femurs were isolated from mice treated with niraparib 25 mg/kg or vehicle and fixed using 4% [w/v] paraformaldehyde (Millipore Sigma) in PBS for 24 hours, followed by a daily increasing sucrose gradient (10 to 30%). Femurs were then frozen using a layer of chilled ethanol and stored at -80°C after embedding in SCEM (R031006, SECTION-LAB Co., Japan) or OCT (23-730-571, Fisher Scientific). Sectioning at ~12 μm thickness was performed on each femur using a tape transfer system[4] at a cryostat (CM3050 S, Leica Biosystems) followed by mounting onto microscopy slides (Fisher Scientific). Cryosections were blocked with 5% goat serum in PBS, followed by primary antibody labeling overnight at 4°C for CD41 (133902, Biolegend) and/or yH2AX (phospho-histone H2A.X, Ser139, 20E3, Cell Signaling Technology). Secondary staining was done with AlexaFluor®-488 and 647-conjugated antibodies raised against the primary rat or rabbit antibody (Invitrogen) and DAPI was used for nuclear staining. Labeled cryosections were imaged using a Lionheart FX microscope with a Phase objective, 4x Olympus Plan Fluorite NA: 0.13 (Ex: 469/35 nm, Em: 525/39 nm) or a Nikon AXR confocal microscope using a 20x objective. MK quantification was performed using an automated imaging platform (Lionheart, Biotek) by selecting CD41 positive objects >15 µm with a nucleus. For yH2AX quantification, yH2AX foci were counted manually with each discrete focus on a confocal plane regarded as a distinct focus.

#### MK Ploidy Analysis

Bone marrow was isolated from 1 tibia from either niraparib 25 mg/kg or vehicle-treated mice by centrifugation at 2500x*g* for 40 seconds. Lysis of red blood cells was performed using ACK buffer (A104921, Gibco) after filtration using a 100 µm cell strainer. Subsequently, cells were washed with PBS, fixed, and permeabilized with 70% ethanol. Cells were then stained with a FITC-conjugated anti-CD41 antibody (133904, Biolegend) and propidium iodide (P1304-MP, Millipore Sigma) in the presence of RNAse A (EN0531, ThermoFisher Scientific). Ploidy of CD41-positive cells was quantified using flow cytometry (BD Accuri C6 Plus).

#### HSPC and MK quantification by flow cytometry

Bone marrow was isolated from 1 tibia and 1 femur per mouse. Single-cell suspensions were analyzed using a Cytek Aurora spectral flow cytometer four laser system (16-V-14B-10YG-8R).

Hematopoietic stem and progenitor cells (HSPCs) were identified by gating on a live, CD45$^{+}$Lin$^{-}$ cell suspension as described previously[2,3]. Populations were defined as follows: multipotent progenitors (MPP2;CD45$^{+}$Lin$^{-}$Sca-1$^{+}$c-Kit$^{+}$Flt3$^{-}$CD48$^{+}$CD150$^{+}$, MPP3; CD45$^{+}$Lin$^{-}$Sca-1$^{+}$c-Kit$^{+}$Flt3$^{-}$CD48$^{+}$CD150$^{-}$, MPP4; CD45$^{+}$Lin$^{-}$Sca-1$^{+}$c-Kit$^{+}$Flt3$^{+}$), MK progenitors (MkP; CD45$^{+}$Lin$^{-}$Sca-1-cKit$^{+}$CD150$^{+}$CD41$^{+}$), MKs (CD45$^{+}$Lin$^{-}$CD41$^{+}$CD42d$^{+}$), short-term HSCs (ST-HSC; CD45$^{+}$ Lin$^{-}$Sca-1$^{-}$c-Kit$^{+}$Flt3$^{-}$ CD150$^{-}$CD48$^{-}$), HSCs (HSC; CD45$^{+}$Lin$^{-}$Sca-1$^{+}$cKit$^{+}$Flt3$^{-}$ CD150$^{+}$CD48$^{-}$), CD41$^{+}$ HSCs (CD45$^{+}$Lin$^{-}$Sca-1$^{+}$cKit$^{+}$Flt3$^{-}$ CD150$^{+}$CD48$^{-}$CD41$^{+}$).

#### Intracellular yH2AX staining

Cells were fixed and processed using the eBioscience™ Foxp3/ Transcription Factor Staining Buffer Set using manufacturer's instructions and stained with anti- yH2AX (600205; Biolegend). Populations were defined similarly as within the HSPC panel; MPP3-MPP4 subset was combined (MPP3-4; CD45$^{+}$Lin$^{-}$Sca-1$^{+}$c-Kit$^{+}$CD48$^{+}$CD150$^{-}$).

#### Electron Microscopy

Femurs from mice treated for 11 days with either vehicle or niraparib (25 mg/kg) were isolated and cleaned. Femurs were fixed with 1.25% paraformaldehyde, 0.03% picric acid, 2.5% glutaraldehyde in 0.1–M cacodylate buffer (pH 7.4) for at least 1 hour. Samples were decalcified in 0.27 M EDTA (pH 7.4) for 1-3 weeks at 4°C, with the EDTA solution changed daily. Samples were then processed by the Harvard Medical School Electron Microscopy core facility. Small pieces (1-2 mm cubes) of fixed tissue were washed in 0.1M cacodylate buffer and postfixed with 1% Osmium tetroxide (OsO4)/1.5% Potassium ferrocyanide ($KFeCN_6$) for 1 hour, washed in water 2x, 1x in 50 mM Maleate buffer pH 5.15 (MB) and incubated in 1% uranyl acetate in MB for 1 hr followed by 1 wash in MB, 2 washes in water and subsequent dehydration in grades of alcohol

(10 min each; 50%, 70%, 90%, 2x10 min 100%). The samples were then put in propylene oxide for 1 hour and infiltrated overnight in a 1:1 mixture of propylene oxide and TAAB Epon (TAAB Laboratories Equipment Ltd, https://taab.co.uk). The following day the samples were embedded in TAAB Epon and polymerized at 60ºC for 48 hours. Ultrathin sections (50-80 nm) were cut on a Reichert Ultracut-S microtome, picked up on to copper grids stained with lead citrate and examined in a TecnaiG² Spirit BioTWIN and images were recorded with an AMT Nanosprint 43-MKII camera. Samples were imaged at 200x. All chemicals (except the TAAB Epon) are from Electron Microscopy Sciences: www.emsdiasum.com.

Supplementary Figure Legends

**Figure S1. PARP inhibitors differentially affect platelet counts.**
A Sysmex hematology analyzer was used to quantify **A)** Immature platelet fraction (IPF) in vehicle or niraparib treated mice and
**B)** Mean platelet volume (MPV) in vehicle or niraparib treated mice (50 mg/kg); n=4-5 mice/group; Data as mean ± SD, Two-way ANOVA with Tukey's correction for multiple comparisons
**C)** Hemoglobin (Hgb) in vehicle or niraparib treated mice (25 mg/kg) and
**D)** White blood cell (WBC) counts in vehicle or niraparib treated mice (25 mg/kg); n=5 mice/ group. Data as mean ± SD, Two-way ANOVA with Tukey's correction for multiple comparisons
Mice were treated with 50 mg/kg olaparib or vehicle as described and
**E)** Platelet counts in vehicle or niraparib treated mice (25 mg/kg).
**F)** Fold change (%) in platelet count from D0 were quantified on indicated days*;* n=5 mice/ timepoint. Data as mean ± SD, Two-way ANOVA with Tukey's correction for multiple comparisons.
Flow cytometry was used to measure **G)** MFI of αIIb integrins (CD41), glycoprotein (GP)Ibα and GPVI in platelets in whole blood; n=5 mice/ group. Data as mean ± SD, unpaired, two-tailed student's t-test.

**Figure S2. Flow cytometry gating strategy for HSPC and MK lineage populations**
**A)** Gating strategy for HSPC populations.

**Figure S3. TEM of whole bone marrow reveals normal MK morphology post niraparib treatment**
Representative images of megakaryocytes within the vascular niche in the bone marrow by electron microscopy. Top = vehicle, bottom = niraparib (25 mg/kg), (scale bars = 2 um). Megakaryocyte nucleus (blue asterisk), red blood cells (RBCs, red filled circle) within the vasculature and the endothelial barrier (green arrows).

**Figure S4. PARP inhibitors increase platelet count without impacting platelet function.**
**A)** Time course of platelet counts in vehicle or niraparib treated mice (25 mg/kg). Mice were followed for an additional 47 days after treatment cessation; n=4-5 mice/ group. Data as mean ± SD, Two-way ANOVA with Tukey's correction for multiple comparisons.
**B)** Absolute counts of HSPC populations 47 days after cessation of niraparib or vehicle treatment were quantified by flow cytometry using the panel in Fig S2 ; n=4-5 mice/ group. Data as mean ± SD, unpaired, two-tailed student's t-test.
**C)** Platelet aggregation after stimulation with CRP or Thrombin was measured 47 days after cessation of niraparib or vehicle treatment; n=4-5 mice/ group. Data is mean ± SD, unpaired, two-tailed student's t-test.
**D** MFI of P-selectin and αIIbβ3 integrin on platelets in whole blood 47 days after cessation of niraparib or vehicle treatment; n=4-5 mice/group. Data as mean ± SD, multiple t-tests.
**E)** Ploidy distribution in native MKs from vehicle or niraparib treated mice (25 mg/kg); n=5 mice/ group. Data as mean ± SD, Two-way ANOVA with Šidák correction for multiple comparisons.

**F)** Viability of ex vivo differentiated MKs in the presence of either niraparib or vehicle; n=4/ condition. Data as mean ± SD.


1. Spindler M, Mott K, Schulze H, Bender M. Rapid isolation of mature murine primary megakaryocytes by size exclusion via filtration. *Platelets*. 2023;34:2192289. doi: 10.1080/09537104.2023.2192289
2. Asquith NL, Carminita E, Camacho V, Rodriguez-Romera A, Stegner D, Freire D, Becker IC, Machlus KR, Khan AO, Italiano JE. The bone marrow is the primary site of thrombopoiesis. *Blood*. 2024;143:272-278. doi: 10.1182/blood.2023020895
3. Barrachina MN, Pernes G, Becker IC, Allaeys I, Hirsch TI, Groeneveld DJ, Khan AO, Freire D, Guo K, Carminita E, et al. Efficient megakaryopoiesis and platelet production require phospholipid remodeling and PUFA uptake through CD36. *Nature Cardiovascular Research*. 2023;2:746-763. doi: 10.1038/s44161-023-00305-y
4. Kawamoto T, Shimizu M. A method for preparing 2- to 50-micron-thick fresh-frozen sections of large samples and undecalcified hard tissues. *Histochem Cell Biol*. 2000;113:331-339. doi: 10.1007/s004180000149
